# Machine learning of stem cell identities from single-cell expression data via regulatory network archetypes

**DOI:** 10.1101/208470

**Authors:** Patrick S Stumpf, Ben D MacArthur

## Abstract

The molecular regulatory network underlying stem cell pluripotency has been intensively studied, and we now have a reliable ensemble model for the ‘average’ pluripotent cell. However, evidence of significant cell-to-cell variability suggests that the activity of this network varies within individual stem cells, leading to differential processing of environmental signals and variability in cell fates. Here, we adapt a method originally designed for face recognition to infer regulatory network patterns within individual cells from single-cell expression data. Using this method we identify three distinct network configurations in cultured mouse embryonic stem cells – corresponding to naïve and formative pluripotent states and an early primitive endoderm state – and associate these configurations with particular combinations of regulatory network activity archetypes that govern different aspects of the cell’s response to environmental stimuli, cell cycle status and core information processing circuitry. These results show how variability in cell identities arise naturally from alterations in underlying regulatory network dynamics and demonstrate how methods from machine learning may be used to better understand single cell biology, and the collective dynamics of cell communities.

## Introduction

The pluripotent epiblast exists transiently in the developing embryo and is the founding tissue for all somatic and germ cells in the adult mammalian organism (Boroviak et al., 2014; Gardner and Beddington, 1988). Because of this remarkable ability there has been sustained interest in deciphering the molecular regulatory mechanisms that underpin pluripotency (Li and Belmonte, 2017). From these studies, it has become increasingly clear that the functional state of pluripotency emerges in a complex, and as yet incompletely understood, way from the collective dynamics of underpinning molecular regulatory networks, which involve numerous protein-protein, protein-DNA, epigenetic and signaling interactions (Azuara et al., 2006; Kim et al., 2008; Kunath et al., 2007; Loh et al., 2006; Meshorer et al., 2006; Niwa et al., 1998; Sato et al., 2004).

The nature of the regulatory relationships in these underlying networks have accordingly become a focus of increasing research attention (Dunn et al., 2014; Xu et al., 2014). Typically, regulatory interactions are inferred from measurements taken from cellular aggregates, usually containing many thousands of cells, and therefore provide an ensemble view that characterizes those interactions that are typical for the ‘average’ pluripotent cell (Gerstein et al., 2012). These ensemble models have been tremendously useful in dissecting the molecular basis of pluripotency and have become successively refined in recent years (Kim et al., 2008; Loh et al., 2006) to include, for example, the processing logic of combinatorial interactions (Dunn et al., 2014; Xu et al., 2014).

However, although undoubtedly powerful tools to understand pluripotency, these networks are fundamentally derived from bulk cell measurements and there is now a need to better understand how these ensemble models relate to regulatory processes within individual pluripotent cells (Filipczyk et al., 2015; Stumpf et al., 2016; Teschendorff and Enver, 2017; Trott et al., 2012).

The relationship between ensemble and individual cell regulatory networks are particularly relevant to the study of pluripotency for two reasons.

Firstly, it is now well observed that apparently functionally homogeneous pluripotent cells exhibit substantial cell-to-cell variability in gene/protein expression patterns, suggesting that pluripotency as a function is compatible with a variety of different molecular configurations (Guo et al., 2016; Kumar et al., 2014; Singer et al., 2014). This has led to acceptance that there are numerous alternate states of pluripotency – most notably naïve and primed states corresponding to the epiblast of the blastocyst, and the epiblast in the egg cylinder of the mouse post-implantation embryo respectively – each with subtly different developmental potential. Our understanding is such that propagation of these alternate pluripotent states *in vitro* is now routine, using different cocktails of growth factor supplementation (Brons et al., 2007; Chou et al., 2008; Evans and Kaufman, 1981; Martin, 1981; Tesar et al., 2007; Weinberger et al., 2016). Importantly, these distinct populations can each contribute to all principal embryonic lineages in *vitro* and are apparently inter-convertible (Chou et al., 2008; Greber et al., 2010; Guo, Ge et al., 2009), suggesting a remarkable plasticity in the dynamics of the underlying regulatory networks. It seems likely that as our understanding of pluripotency develops, other varieties of pluripotency will be discovered and sustained *in vitro*. Indeed, it has recently been proposed that pluripotent cells also progress through an important *formative* state, in which the naïve regulatory network is partially dissolved and cells become competent for lineage allocation (Kalkan and Smith, 2014; Smith, 2017).

Secondly, the epiblast appears insensitive to the removal or addition of cells (Gardner and Beddington, 1988), suggesting a level of functional redundancy between individual cells that is supportive of the notion that pluripotent cell populations in vivo behave more like a ‘collection of transition cells’ (Gardner and Beddington, 1988), than a defined developmental state *per se*. This collective behavior presumably also emerges from the dynamics of the underlying regulatory networks, although the mechanisms by which such collective dynamics are regulated by intracellular regulatory networks is still largely mysterious (MacArthur and Lemischka, 2013). Taken together, these findings suggest that the regulatory network underlying pluripotency exists in a number of interchangeable configurations, although the nature of these different configurations, and their relationships to one another, are not yet fully understood (Stumpf et al., 2016; Trott et al., 2012).

Here, we sought to develop a method to interpret single cell data to better understand how alterations in regulatory network activity within individual cells gives rise to variability within pluripotent cell populations.

To approach this problem, we were inspired by a method from the early days of face recognition, which de-constructs facial images into facial archetypes, known as *eigenfaces*, that are learned from a training set of portraits, and reconstructs unseen faces as weighted sums of these learned eigenfaces (Sirovich and Kirby, 1987; Turk and Pentland, 1991) (see Fig. 1). Although face recognition methods are now highly sophisticated, the original implementation of the eigenface routine is essentially an ingenious, although mathematically straightforward, implementation of principal component analysis (PCA) that relies on the fact that each facial image may be considered as a matrix of numbers, and therefore reshaped to a vector and associated with a point in a high-dimensional space. Thus, given a set of training portrait images, PCA may be used to extract the characteristic features – the eigenvectors of the training covariance matrix, also known as principal components – that capture significant variation within the training set (Fig. 1a). By transforming these eigenvectors back into matrices of the same dimension as the images in the training set they can be visualized as facial archetypes (or ‘eigenfaces’) of the training set (Fig. 1a). Remarkably, it was observed that only a small number of eigenfaces (typically ∼ 5%) is sufficient to explain 95% of facial details, and therefore unseen portrait images can be reliably reconstructed as a weighted sum of a very small number of eigenfaces (Fig. 1a). Importantly, this means that the vector of weights alone (i.e. ∼5 numbers) is typically sufficient to recognize an individual from their portrait, thus significantly reducing the dimension of the recognition problem (Fig. 1b).

**Figure 1:**
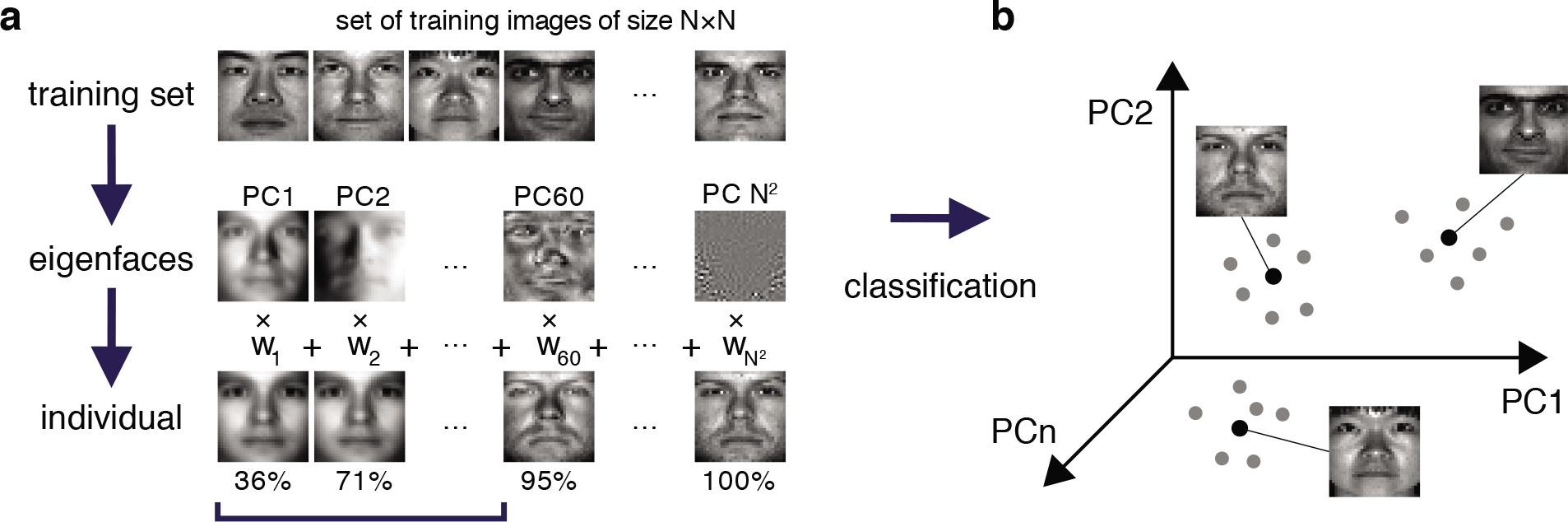
Eigenfaces for face recognition. **(a)** A training set of portrait images of size *N* × *N* is used to extract the facial archetypes (eigenfaces) encoded by the *N*^2^ principal components of the training set. A small subset of eigenfaces explains most of the variability in facial features between individuals. In this specific example from The Extended Yale Face Database B (Georghiades et al., 2001; Lee et al., 2005), a recognizable version of an original test image can typically be reconstructed from a weighted sum of the first 5:9% (60 out of 1024) eigenfaces, which explain 95% of the variance in the data. **(b)** Each test face may be reconstructed as a weighted sum of eigenfaces, and thereby efficiently encoded by a weight vector, which may be thought of as a point in a much lower dimensional space than the original feature space. In this case although each face is initially associated with a point in a 1024 dimensional space (corresponding to the 1024 pixels in the original image), a recognizable version may be reconstructed in just 60 dimensions (the corresponding weightings). Different images of the same person typically occupy a region in the principal component space around a central characteristic image.

While this does not immediately appear to relate to the study of pluripotency we surmised that a similar approach could be used to reconstruct pluripotent cell identities from single cell data, as a weighted sum of regulatory network archetypes. Furthermore, just as portrait images can be efficiently encoded using the eigenface weight-vector, we wanted to determine if complex patterns of gene/protein expression within individual cells could similarly be encoded by a low-dimensional representation in terms of the activity of these network archetypes, thereby facilitating more accurate classification of cell identities from noisy expression data.

## Results

### Integrating regulatory interactions with single cell data

We first sought to obtain a reliable training dataset of protein expression patterns in pluripotent cells across multiple intracellular information levels, including the protein abundance of core transcription factors (Kim et al., 2008; Loh et al., 2006), the phosphorylation status of signaling pathways (Kunath et al., 2007; Niwa et al., 1998; Sato et al., 2004) and global transcriptional activity based on histone acetylation (Azuara et al., 2006; Meshorer et al., 2006). Such systems-level proteomic information at single-cell resolution is currently only available through immunolabeling followed by mass-cytometry, a highly specialized technique that is available to only a small number of groups (Spitzer and Nolan, 2016). Thus, we sourced a relevant training dataset from the literature (Zunder et al., 2015). In total this training data consists of expression patterns of 34 proteins and protein modifications in 31,876 pluripotent cells from two mouse embryonic stem cell (mESC) lines (Nanog-GFP [NG] mESCs and Nanog-Neo [NN] mESCs that express green fluorescent protein [GFP] or a Neomycin resistance gene respectively from the endogenous Nanog locus (Wernig et al., 2008)), grown in low-serum medium supplemented with Leukemia Inhibitory Factor (LIF; 0i conditions). In addition, this dataset also contains expression levels of the same features in 15,540 NG mESCs and 15,752 NN mESCs grown in medium supplemented further with a GSK3β inhibitor and a MEK inhibitor (known as 2i conditions, which support the pluripotent ‘ground’ state (Ying et al., 2008)), as well as expression time-course data containing 834,548 secondary mouse embryonic fibroblasts (MEFs) generated from both cell lines that express Yamanaka reprogramming factors (Takahashi and Yamanaka, 2006) under the control of a doxycycline (dox) inducible promoter (Wernig et al., 2008).

To interrogate this data, we sought to supplement it by constructing a directed regulatory network specific to the features (transcription factors, surface epitopes, phosphorylation, etc.) that had been quantified (Fig. 2). Features (that is, proteins profiled) in this signed, directed regulatory network are represented as nodes and regulatory interactions between features are represented as edges between pairs of nodes (an edge is positive if it is activating, and negative if it is inhibiting). Evidence for node interactions was extracted from transcription factor binding data from ChIPBase 2.0 (Zhou et al., 2017), and information on other known interactions were sourced from the Kyoto Encyclopedia of Genes and Genomes (KEGG) (Ogata et al., 1999) and Reactome (Fabregat et al., 2016) (see Table S1 for details). Unconnected nodes, such as the inert GFP reporter, and cell cycle markers pH3 and IdU were removed from the analysis. The resulting network G contains 27 nodes, connected by 124 edges (Fig. 2a).

**Figure 2:**
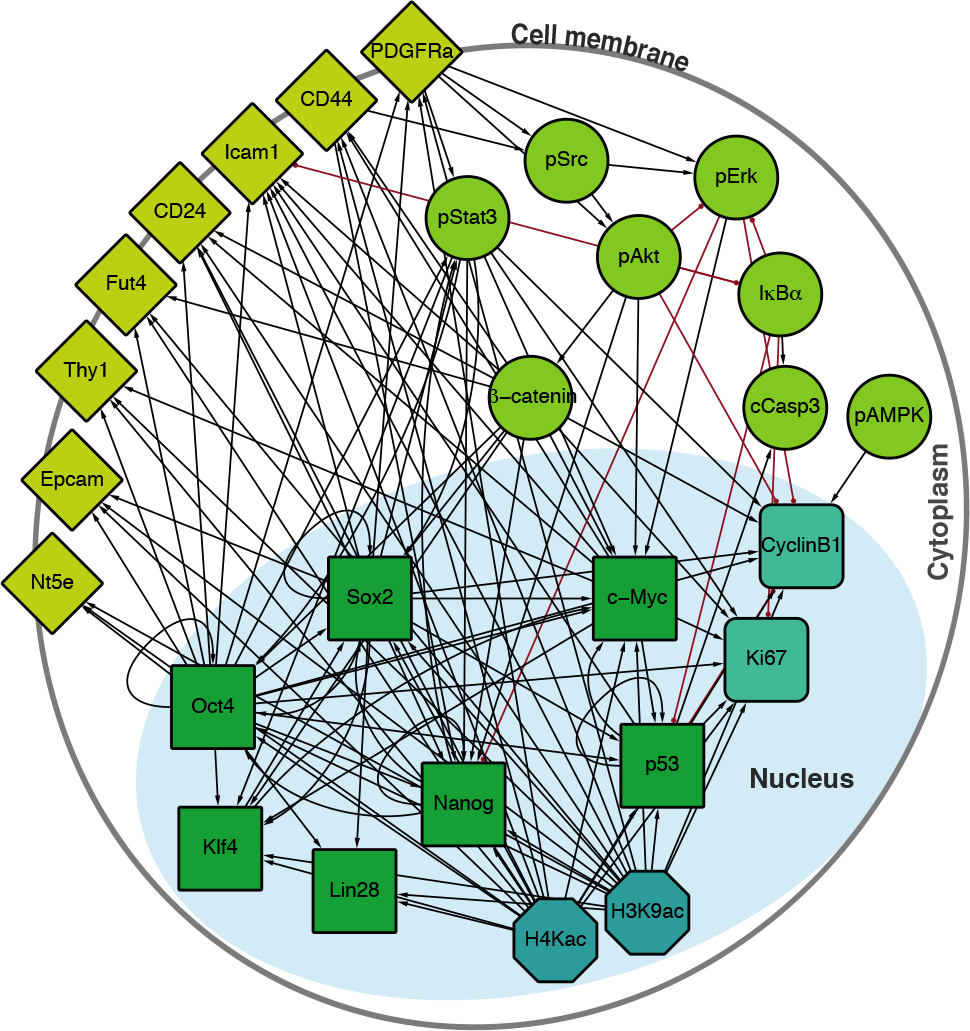
Integrated regulatory network derived from the literature. Schematic shows the structure of the inferred regulatory network between the factors profiled, derived from the literature (see Table S1). The network accounts for multiple molecular information processing mechanisms, at multiple different spatial locations in the cell, including interactions between: transcriptional regulators (green squares), chromatin modifiers (petrol octagons), cell cycle factors (sea green rounded squares), signaling cascades (light green circles), and surface molecules (yellow diamonds).

The overall structure of *G* is conveniently encoded in the network adjacency matrix,

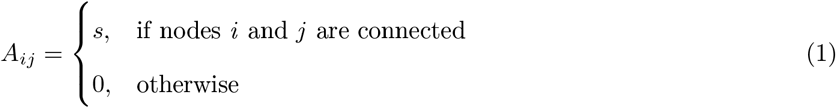

where *s* = +1 for activating interactions, and *s* = −1 for inhibitory interactions.

The first step in our process consists of combining this regulatory network with the single cell expression training set. Trivially, the expression data represents the activity of the nodes in the network within each cell, but does not take into account regulatory interactions between nodes. To incorporate this information, we assumed that the activity of each edge within the network is determined by the signal intensities of both interaction partners within the individual cell. Accordingly, denoting the vector of expression values in a given cell by ***v***, we created a weighted adjacency matrix ***W***

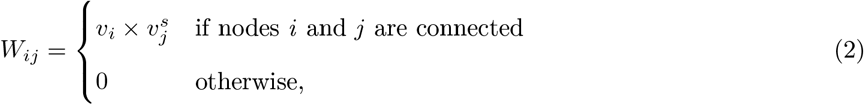

where the sign of an edge *s* ∈ [−1, +1] denotes either inhibiting or activating interactions. Thus, we associated a high weight to a positive edge if both the source and the target were highly expressed, and a high weight to a negative edge if the source was highly expressed and the target was expressed at a low level. Informally, this representation may be thought of as assigning high confidence that a given edge is expressed within an individual cell if its source and target nodes are expressed consistently with the sign of the edge relating them. The resulting weighted adjacency matrix ***W*** is a simple measure of the extent to which the network *G* is expressed in the cell given the expression patterns observed in that cell. By analogy with the face recognition problem, ***W*** may be considered as the ‘image’ of the cell.

As with the eigenface routine, this matrix may be easily restructured as a vector. In this case, ***W*** may be coerced into a vector of length *m* (where *m* is the number of edges in the network, here 124), by first reshaping it to a vector of length *n*^2^ (where *n* is the number of nodes in the network, here 27), and then squeezing out all entries for which *A_ij_* = 0. This procedure effectively injects the expression data with prior knowledge of the network structure, leading to an expansion of the original feature space from ℝ^*n*^ to ℝ^*m*^ (generically a connected network will have more edges than nodes, unless it is a tree). Using this method, we inferred the activity of the regulatory network *G* within each of the ∼ 9 × 10^5^ individual cells profiled. For subsequent analysis we treated NG mESCs cultured in 0i conditions as a training dataset and held back the remaining data to test the model learned from the training data.

### Regulatory networks characterize alternate states of pluripotency

Once the training data had been produced, we conducted principal component analysis. In the same way that the principal components (PCs) in the eigenface routine may be reshaped and interpreted as facial archetypes from which individual portraits may be reconstructed, the principal components here may be reshaped and interpreted as network archetypes from which pluripotent cell identities may be reconstructed. However, while only ∼ 5% of the PCs are required for accurate face recognition, we found that (for both NG and NN mESCs) ∼ 23% of the PCs were required to explain 95% of the variance in our training data (Fig. 3a). The larger number of PCs required is not unexpected, and is reflective of the high levels of noise that are characteristic of high-throughput single cell data (Graf and Stadtfeld, 2008). Therefore, rather than using the proportion of variance explained to determine the appropriate number of PCs to retain for subsequent analysis, we sought to identify the minimal number needed to preserve the natural clustering structure in the data.

**Figure 3:**
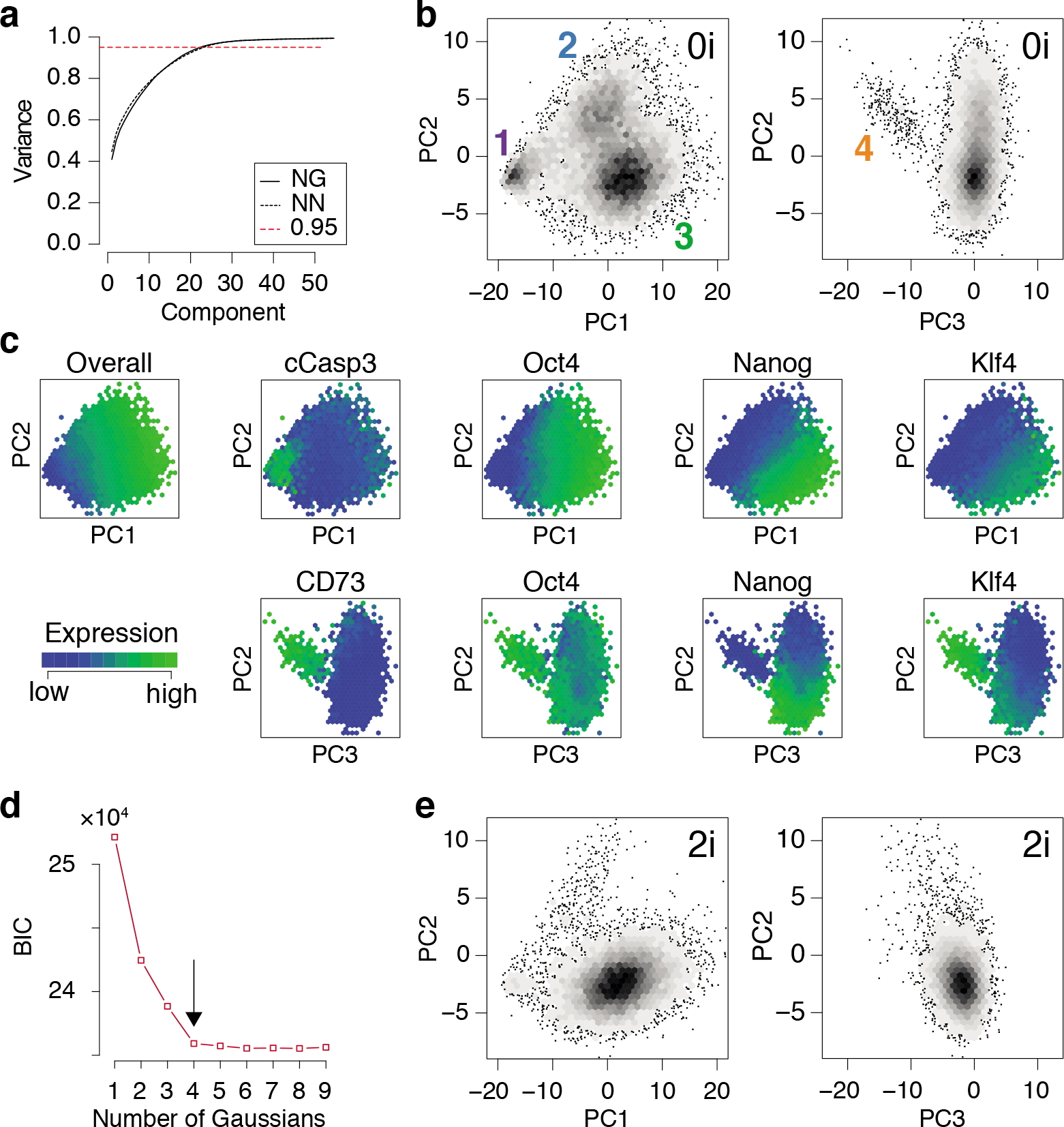
PCA identifies three distinct pluripotent states within ES cells cultured in 0i conditions. **(a)** Cumulative proportion of variance explained by principal components (PCs) of the training data for Nanog-GFP (NG) mESCs and Nanog-Neo (NN) mESCs respectively. The dotted red line marks the commonly used threshold value of 0:95. **(b)** Density plot of training data from NG mESCs projected onto the first three components. Four clear clusters are apparent, labeled 1-4, corresponding to distinct states of network activity. Each hexagonal bin contains at least 5 cells. **(c)** Heat map of expression of important nodes in NG mESCs projected onto PCs 1-3. Mean expression values are displayed for each hexagonal bin. Distinct alternate states of pluripotency are apparent, based upon edge co-expression patterns. **(d)** Bayes information criterion (BIC) as a function of the number of Gaussian mixture components fitted to the first three principal components. The arrow marks the elbow in the plot, indicating the optimal number of components (here 4). (e) Projection of a test dataset of expression patterns from NG mESCs cultured in 2i conditions onto the training PCs (panel b). Panels b-e show data from NG mESCs, corresponding data for NN mESCs is shown in Fig. S1.

We found that four distinct clusters of cells were readily identifiable in the full dataset (natural clustering structure was obtained by fitting a Gaussian mixture model to the data and selecting the model that minimizes the Bayesian information criterion [BIC], see Fig. 3d and Fig. S1d). This natural clustering was robustly retained when projecting the data onto the first three PCs (Fig. 3b); higher components only added noise to this basic clustering structure. This analysis suggests that PCs 1-3 account for the biological variability present in the data, while higher components primarily correspond to technical variability.

Since the PCs are linear combinations of the underlying features (here, network edges) each one may be thought of as regulatory network archetype, and the expression pattern of each cell in the training data may therefore be reconstructed as a weighted sum of these archetypes. By analogy with eigenface routine, we will call these network archetypes *eigen-networks*. Since PCs 1-3 account for the biological variability in the data, the structure of the eigen-networks associated with these components are of particular interest. The first eigen-network (PC1 in Fig. 4a) naturally separated cells into two subsets (Fig. 3b), based upon overall activity of regulatory interactions (see Fig. 4a and overall expression in Fig. 3c). A subset of cells with low overall edge expression (cluster 1 in Fig. 3b) primarily contained apoptotic cleaved Casp3-positive cells (Fig. 3c) and cell cycle arrested cells (Fig. S1g), likely caused by the increased activity of IκBα (Fig. 4a). Cluster 1 also lacked activity between the core pluripotency factors Oct4, Nanog and Klf4 (Fig. 4a and Fig. 3c). In contrast to this small subset, the majority of cells displayed high overall expression of pluripotency related-factors, including Oct4 (compare positive edge association in PC1 Fig. 4a and node expression in Fig. 3c).

**Figure 4:**
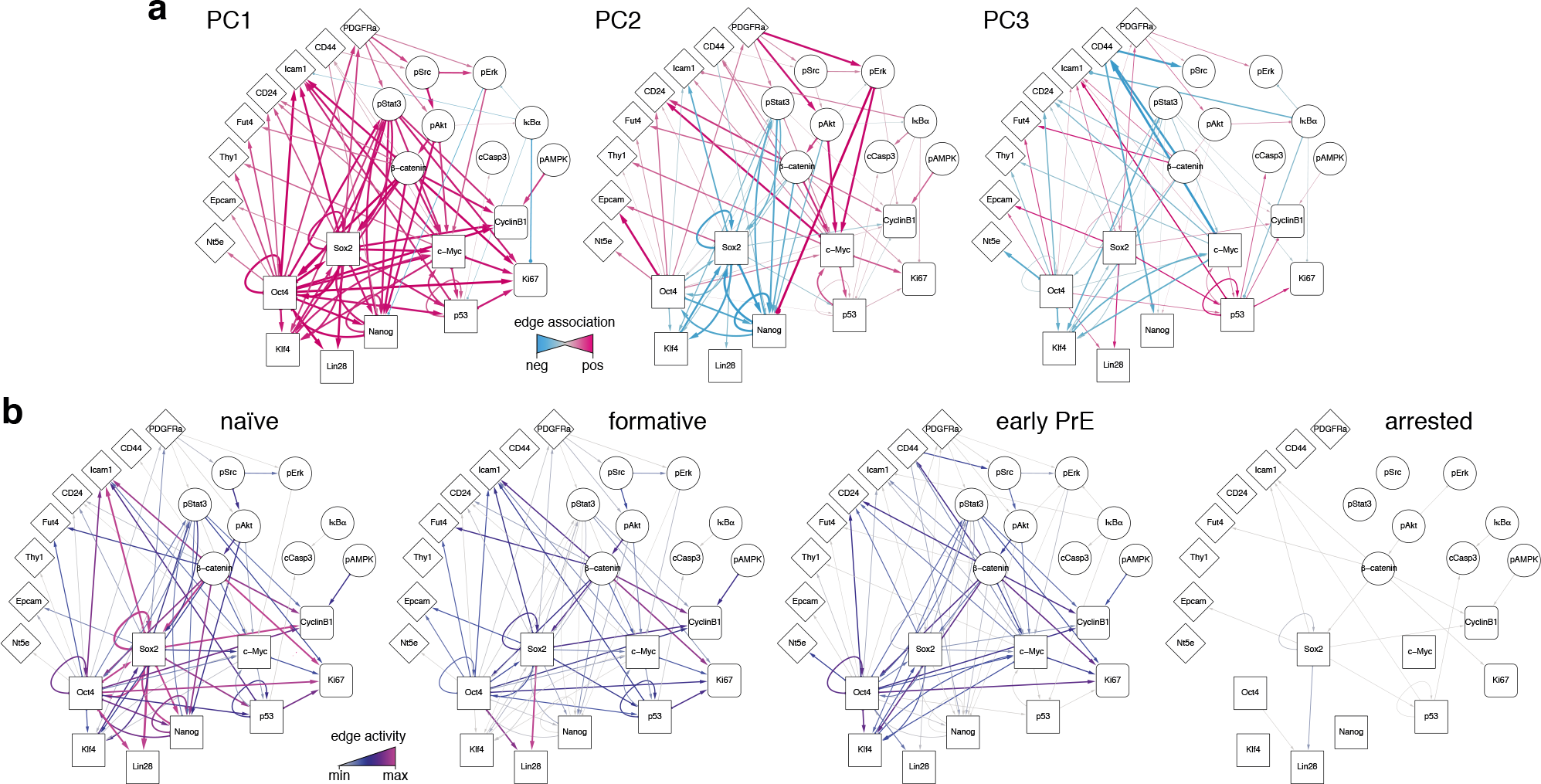
Regulatory network activity archetypes define alternate pluripotency states. **(a)** Graphical representation of the first three PCs, interpreted as regulatory network archetypes. Color and edge width indicate signed deviation from the mean. **(b)** Representative regulatory network states for naïve, formative, and early primitive endoderm (PrE) states. The network corresponding to arrested/apoptotic cells (cluster 1) in Fig. 3b is also shown for reference.

The majority pluripotent population identified by the first eigen-network naturally separated into 2 distinct further sub-populations (clusters 2, 3 in Fig. 3b) by expression of the second eigen-network (PC2 in Fig. 4a), which broadly captures the strength of connection between the cell’s signaling pathway activity and its core transcriptional regulatory circuitry, including activity of *β*-catenin (Wnt-signaling), Stat3-phosphorylation (LIF-signaling) and Erk-phosphorylation (FGF/MEK-signaling) (blue edges in Fig. 4a, PC2). This component therefore captures integration of the primary axes of extrinsic control of the pluripotent ground state (Ying et al., 2008), and distinguishes cells in the pluripotent ground state (cluster 3), which are characterized by high Nanog, Oct4 and Klf4 expression and strong integration of signaling and core transcriptional regulatory programs, from those in a second pluripotent state (cluster 2), which are characterized by low Nanog and Klf4 expression (Fig. 3c), and more sporadic connectivity between signaling and transcriptional controls and high Erk-signaling activity (red edges in Fig. 4a, PC2). This expression pattern indicates that these cells may correspond to a more developmentally advanced state (Marks et al., 2012). While the full nature of this state has yet to be determined, it is consistent with the recently proposed ‘formative’ phase of pluripotency, characterized by dissolution of core pluripotency sustaining mechanisms (Smith, 2017).

In addition to these primary populations we also observed small subset of cells (∼ 2%) that could be distinguished from the formative and naïve pluripotent states based on expression of the third eigen-network (see population 4 in Fig. 3b). This fourth population is similar to the formative state (population 2) with respect to expression of Nanog (both low; see Fig. 3c) and similar to the naïve state (population 3) with respect to expression of Klf4 (both high; see Fig. 3c). However, it is quite distinct with respect to a number of surface markers. Notably cells in cluster 4 are CD73^high^ (Nt5e; Fig. 3c), and CD44^high^ and CD54^low^ (Fig. 4, PC3), suggesting an increased interaction with the extracellular matrix. These differences are not simply a manifestation of mitosis or cell cycle arrest, since the proportion of M-phase cells in this population is comparable to both the naïve and formative states and the proportion of G0-phase cells is comparable to the formative state (Fig. S1f-g). Although this data does not include more specific markers such as Gata6 and Sox17, we conjecture that this population corresponds to the early primitive endoderm (PrE), due to the observed low expression of Nanog and co-expression of Oct4 and Klf4 (Boroviak et al., 2014; Guo et al., 2010). Additionally, these cells display high levels of STAT3 signalling acitvity (blue edges in Fig. 4a, PC3), which has been shown to support PrE differentiation (Morgani and Brickman, 2015). Moreover, in the process of PrE differentiation, cells undergo an epithelial-to-mesenchymal transition (EMT) (Chazaud et al., 2006) and begin to express mesenchymal markers such as CD73 (see Fig. 3c and the blue edge between Oct4 and Nt5e, and between Oct4 and CD24 in Fig. 4a, PC3). In accordance with this notion, we observe that this population has the highest total within cluster variance, indicating the presence of substantial cell-cell variation (see Fig. S1e), which is typically found in cells transitioning from one state to another (Bargaje et al., 2017).

To investigate this possibility further we constructed representative networks for each of the four identified states using the first three eigen-networks and the weight vector corresponding to the centroid for each cluster (see Fig. 4b). The resulting networks may be thought of as representations of the characteristic patterns of network activity within each of the four states we identified. These networks show that: (1) the pluripotent ground state is characterized by strong co-regulatory activity between members of the core transcriptional circuit and strong integration of signalling pathways with this core sub-network (Fig. 4b). (2) By contrast, the PrE state is characterized by partial dissolution of the core transcriptional circuit (in particular a loss of Nanog, Sox2 and p53 activity), which is accompanied by changes in cell-cell (CD54) and cell-matrix (CD73, CD44) mediated signaling. However, cells in this state continue to perceive environmental signals via the LIF/Stat3 signaling pathway (Fig. 4b), indicating continued receptivity to pluripotency-stimulating environmental cues. (3) The putative formative state is marked by a further dissolution of the core transcriptional circuit, including the loss of Klf4 regulatory activity (Fig. 4b) and a decrease in LIF/Stat3 signaling (Fig. 4b), suggesting that these cells are transitioning away from the pluripotent ground state. Accordingly, the formative state is also marked by the positive regulation of EpCAM (Fig. 4b), suggesting the onset of cell polarization, as is observed in the epiblast of the egg cylinder *in vivo* (Bedzhov and Zernicka-Goetz, 2014).

In summary, this analysis revealed the presence of four distinct cellular communities, each characterized by different levels of activity of regulatory network archetypes, within mouse ES cell populations cultured in 0i conditions. To determine how general these results were we also examined network expression patterns mESCs cultured in 2i conditions, which stimulate Wnt signaling activity and reduce Erk-phosphorylation using small molecule inhibitors of MEK, and thereby shield the core transcriptional circuitry from extrinsic differentiation cues (Ying et al., 2008). In accordance with the nature of these conditions we found that populations 1, 2 and 4 (corresponding to arrested, formative and PrE cells) were comprehensively depleted in mESCs cultured in 2i conditions, while cluster 3 (corresponding to the naïve or ground state) was robustly maintained (Fig. 3e).

These results re-affirm the potency of these conditions to purify the ground state of pluripotency, and provide mechanistic insight into the molecular mode of action of these conditions.

### Individual cells transition through distinct network activity states during reprogramming

To further investigate the biological importance of the regulatory network archetypes we had identified we then sought to determine their temporal expression during cellular reprogramming of somatic cells to pluripotency.

During cellular reprogramming, pluripotency regulatory network activity is typically initially established through the ectopic expression of four trans-genes, Oct4, Sox2, Klf4 and c-Myc (OSKM) (Takahashi and Yamanaka, 2006). Subsequently, the concerted action of these core reprogramming factors leads to profound changes to the cellular phenotype, ultimately re-instating a self-sustaining pluripotent identity in a small proportion of cells. The dynamics of this process are thought to be initially driven by low frequency stochastic events followed by the deterministic progression through a series of characteristic intermediate, partially reprogrammed, expression states (Buganim et al., 2012). It is presumed that these intermediate partially reprogrammed states correspond to partial re-configurations of the pluripotency regulatory network (Golipour et al., 2012). However, the relationships between regulatory network reconfigurations and the dynamics of reprogramming are not well understood.

To address this issue, we considered data from a reprogramming time-course in which the expression of ectopic OSKM transgenes were induced in secondary MEFs by doxycycline (dox) supplementation of the MEF culture medium for 16 days, followed by a further 14 days in 0i conditions without dox (Zunder et al., 2015).

To analyze this data we first fit our training data (expression patterns of NG mESCs cultured in 0i conditions) projected onto the first three eigen-networks (as described above) with a Gaussian mixture model (GMM) with four components. This GMM may be thought of as an estimate of the joint probability density function *P*(*x*) for the training data, projected onto the first three PCs (where *x* ∈ ℝ^3^ identifies points in PC space). We then projected the reprogramming time-course data onto the first three PCs derived from the training data and used the fitted GMM to estimate the likelihood of observing the expression patterns seen in the reprogramming time-course within the pluripotent cell population. That is, if *v* is the expression pattern of a given cell in the reprogramming time-course projected onto PCs 1-3 from the training data, we calculated *P*(*v*) as a measure of the likelihood of observing *v* in the training population. The negative logarithm of this probability

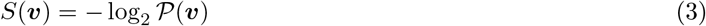

is the amount of information imparted by observation *v* with respect to the probability measure *P* (Cover and Thomas, 1991). Informally, *S*(*v*) is a measure of the ‘surprisal’ of observing the expression pattern *v* in a pluripotent population: cells that express proteins in a pattern similar to that often seen in pluripotent cells have a low surprisal; while cells that express proteins in a pattern that is unusual for pluripotent cells have a high surprisal. To obtain assessment of the dynamics of reprogramming, we calculated the surprisal for each of the 263,692 NG cells in the reprogramming time-course, and monitored how the distribution of surprisal in the population changed over time during reprogramming.

We first observed that the surprisal remained high, and approximately constant, for the first 10-12 days of reprogramming (Fig. 5a), indicating that cells in the starting population (in this case NG MEFs) consistently exhibited expression patterns that are unusual for pluripotent cells, as expected. However, around days 10-12 the population split into two distinct sub-populations: a majority sub-population in which the surprisal remained high, and a minority sub-population in which the surprisal was substantially reduced, suggesting the emergence of population of pioneer partially reprogrammed cells (Fig. 5a). Over the next approximately 20 days the proportion of cells in the low surprisal sub-population gradually increased, indicating the consolidation and proliferation of a robustly pluripotent population of cells (Fig. 5a).

**Figure 5:**
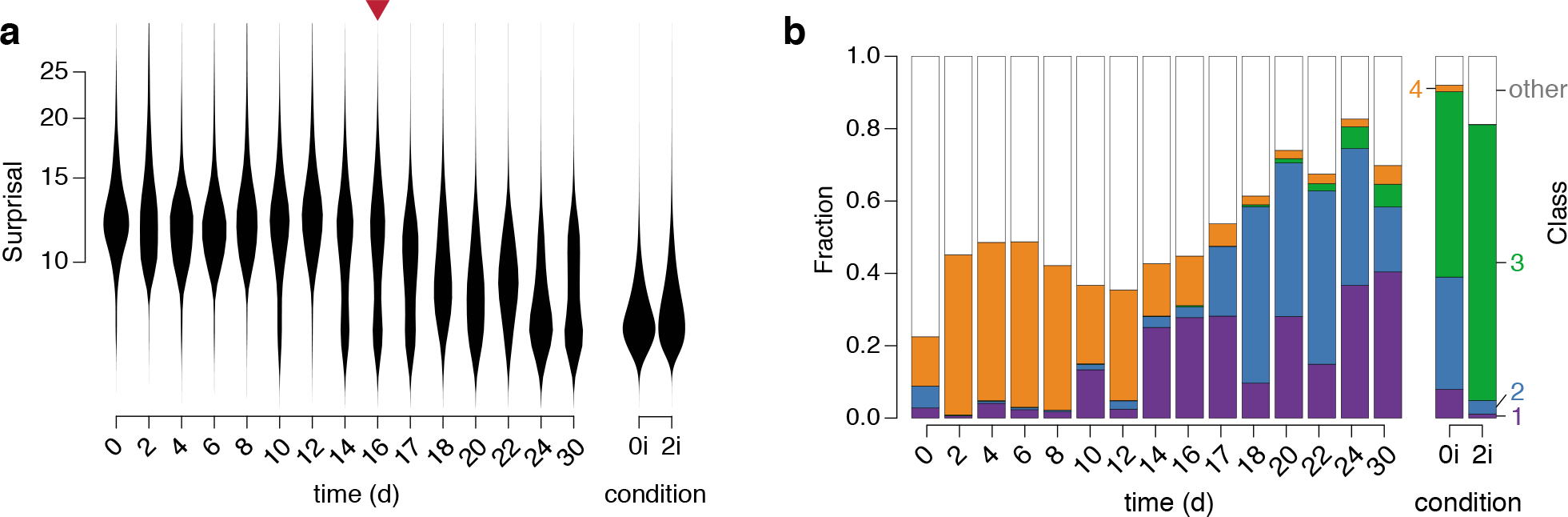
Dynamics of regulatory network activity during cellular reprogramming. **(a)** Violin plots of changes in ‘surprisal’ (Eq. (3)) over time. A gradual decrease in surprisal in the population accompanies cellular reprogramming. The red arrow marks the end of doxycycline treatment. **(b)** The fraction of cells classified into each of the four clusters identified in the training data. Class labels are as in Fig. 3b.

To better understand the identity of this emerging pluripotent sub-population we sought to relate it to the three alternate pluripotency states we had identified (see Fig. 5). To do so we used our fitted GMM to classify each cell in the time-course into one of the four populations identified in the training data (Fig. 5b). Since numerous cells, particularly at the beginning of the time-course, did not resolve well onto any of the clusters in the training data (which is to be expected, since they are not pluripotent) we also incorporated a fifth class to capture those cells with network activity states that were distinct from those found in the training data (for details see Methods).

This analysis revealed that specific instances of regulatory network activity define distinct phases of the reprogramming process (Fig. 5b).

Initially, while the majority of cells were unclassified, indicating lack of similarity to all of the pluripotent training populations, a small proportion of cells were associated with the fourth cluster, corresponding to the early PrE in 0i conditions. This observation is not unexpected as these early PrE cells express Oct4 and Klf4 in addition to surface markers CD24, CD44 and CD73 (see Fig. 3c). Similarly, in the presence of dox, MEFs initially express exogenous OSKM transgenes in parallel to endogenous mesenchymal surface markers such as CD44 and CD73 that are normally expressed in MEFs, until undergoing the mesenchymal-to-epithelial transition (Li et al., 2010). Therefore, these cells display regulatory configuration similar to the early PrE state. This route is consistent with previously observed expression sequence of CD44, Icam1 and Nanog during reprogramming (O’Malley et al., 2013).

This initial phase is followed by the emergence of a population of cells in cluster 1 (corresponding to arrested or apoptotic cells that are frequently observed in reprogramming (Smith et al., 2010)) from day 10-14, followed closely by the emergence of a population of cells in cluster 2 (corresponding to the formative pluripotent state) from day 17 and lastly, the emergence of a small population of fully reprogrammed cells in cluster 3 (corresponding to the pluripotent ground state) after 22 days.

These data suggest that reprogrammed cells do not emerge in significant numbers until after after dox is withdrawn, at which point the regulatory network begins to assume a more natural configuration similar to that of the formative state. These observations are in accordance with the notion that activation of the OSKM transgenes prevent cells from entering a stabilization phase of reprogramming in which the pluripotent state becomes fully established (Golipour et al., 2012). Notably, at around the same time there is an apparent reduction in the frequency of cluster 4 cells, which are marked by low Sox2 and p53 activity, indicating that these cells only exist transiently during reprogramming. Since this population is more variable than the naïve and formative pluripotent populations, it may also mark the handover from the early stochastic phase of reprogramming, in which the activation of OSKM transgenes initiate transformation of the regulatory network configuration, to the late deterministic phase, in which the pluripotent cell identities are consolidated by endogenous regulatory mechanisms (Buganim et al., 2012).

Taken together these results indicate that reprogrammed MEF cells enter pluripotency via a PrE-like state. It remains to be seen if this is a general characteristic of reprogramming that also applies to cells of different somatic origin, or if this particular route is due to the fact that the MEF starting population has a mesenchymal origin that happens to be more similar to the PrE state than it is to the other pluripotent identities (see Fig. 6). Indeed, it was recently demonstrated that reprogramming with the OSKM cocktail can also result in induced extra embryonic endoderm (iXEN) stem cells in parallel to fully reprogrammed iPSCs (Parenti et al., 2016).

**Figure 6:**
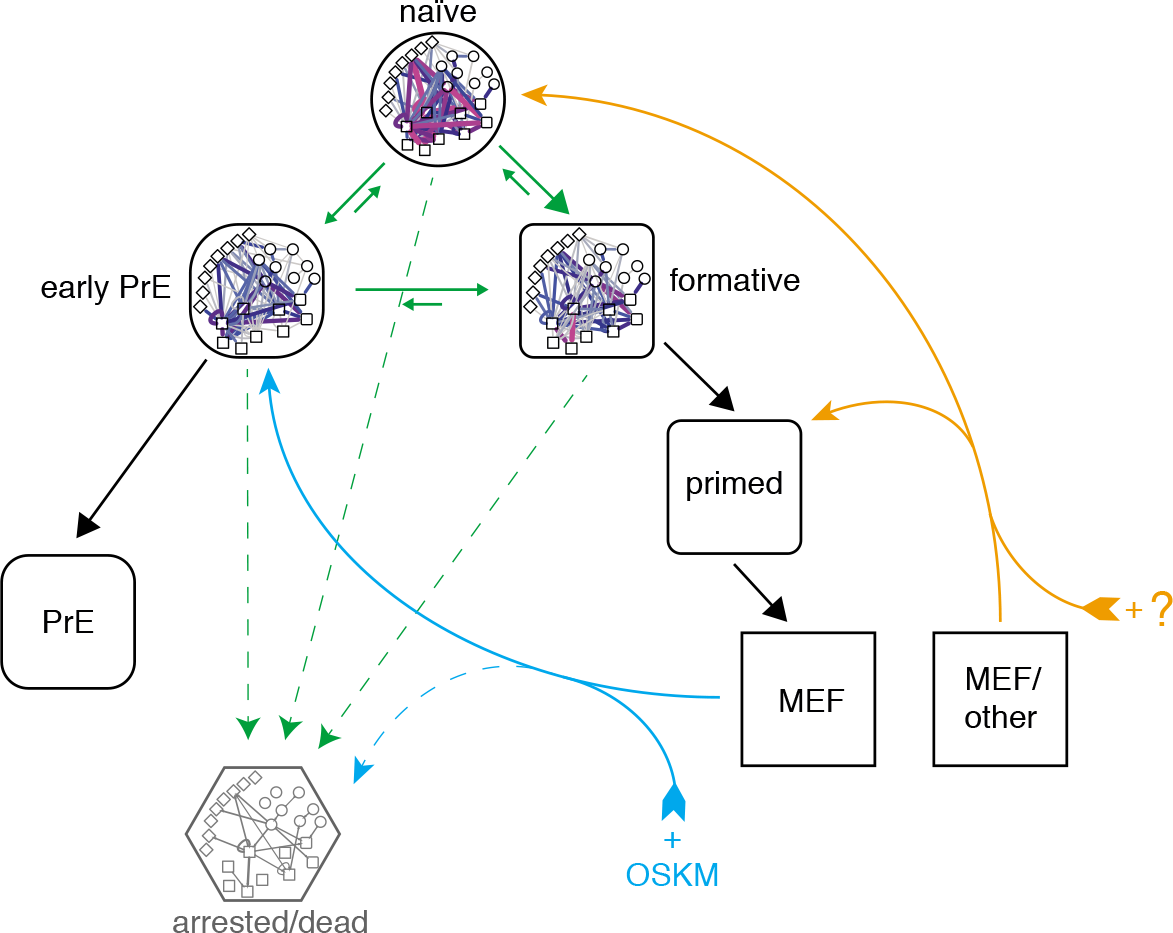
Proposed topography of pluripotency states. Cells descend a natural hierarchy of distinct regulatory network activity states (black arrows). Despite this natural hierarchy, 0i culture conditions permit the inter-conversion of these different network configurations *in vitro* (green arrows). Somatic cell reprogramming re-establishes a network configuration similar to that of the early primitive endoderm (PrE; cyan arrows), from which cells replenish the remaining pluripotency states. A different cocktail of reprogramming factors may enable different reprogramming trajectories to the formative or naïve states from different starting populations (orange arrows). A subset of cells also undergo cell cycle arrest or apoptosis (dashed arrows).

Although the approach taken in this study is centered on pluripotent network configurations observed in steady-state culture conditions, our analysis of network reconfiguration dynamics during reprogramming is consistent with the detailed clustering performed by Zunder et al., who report that cells initially transition through a Oct4^high^/Klf4^high^ state and increasingly resemble partially reprogrammed, transgene-dependent cells prior to mesenchymal-to-epithelial transition (MET) (Zunder et al., 2015). Based on the similarity of these partially-reprogrammed cells with the PrE-like network state we have identified, and the fact that MET provides a major obstacle in somatic cell reprogramming (Li et al., 2010), we propose that futher study of PrE commitment may also help understand the late phase of cellular reprogramming.

## Discussion

The notion that there is a single well-defined pluripotent stem cell identity has been rapidly eroded by advances in single cell analysis methods, which are now revealing ever greater varieties of pluripotency (Guo et al., 2016; Kumar et al., 2014; Singer et al., 2014; Ying et al., 2008). Collectively, these results suggest that pluripotency is not a single phenotype but instead is a property that spans a continuum of observable cell states (Gardner and Beddington, 1988; Morgani et al., 2017; Silva and Smith, 2008; Smith, 2017; Stumpf et al., 2017; Ying et al.,2008). This is in part because the densely connected pluripotency regulatory network is rich in feedback loops which both stabilize pluripotency, and endow pluripotent cells with a remarkable phenotypic plasticity (Kim et al., 2008; MacArthur et al., 2012). Hence, to fully understand pluripotency, strategies to decipher regulatory networks at single cell resolution are needed.

There have been a number of notable advances to this end, particularly with regard to methods for inferring and analyzing regulatory networks directly from single cell data, which can reveal aspects of regulatory control that are inaccessible to study with ensemble techniques such as ChIP-Seq (Buganim et al., 2012; Chan et al., 2017; Stumpf et al., 2017; Trott et al., 2012). For example, Trott and co-workers have inferred regulatory network activity from correlation patterns in single cell data in different stem cell sub-populations, and related these different activity patterns to different aspects of the stem cell identity (Trott et al., 2012). Similarly, Stumpf (not the current author) and colleagues have used powerful notions from information theory to more precisely identify regulatory interactions from single cell time-course data (Chan et al., 2017). However, single cell data is inherently noisy, and consequently large numbers of cells are needed to gain the statistical power to accurately distinguish functional from spurious interactions (Chan et al., 2017).

To circumvent this problem here we have presented a method that incorporates prior knowledge of regulatory interactions directly into single cell expression patterns, rather than inferring regulatory interactions from the data itself, and uses this prior knowledge to dissect the regulatory processes that give rise to different states of pluripotency. This approach is similar to that taken by Teschendorff and colleagues, who, by projecting single cell data onto a known regulatory network, find that pluripotency can be remarkably well related to systems-level emergent network properties (Teschendorff and Enver, 2017). We anticipate that as single cell profiling methods develop we will see concurrent advances in the statistical methods needed to investigate and interrogate the resulting data: indeed, new statistical advances will be essential to fully realize the power of these new and emerging technologies. We expect that Bayesian methods, which use known regulatory interactions as a prior to guide learning of functional interactions directly from single cell data, will combine the benefits of the two approaches to this problem and may therefore be particularly powerful.

In summary, we have adapted a simple image analysis method to infer the presence of four distinct patterns of pluripotency, based on the activity patterns of three regulatory network archetypes within individual cells. The power of our method is not due to its mathematical or computational sophistication – indeed, it is mathematically and computationally straightforward – but rather in the biological interpretation it allows. As such it provides a simple example of how methods from machine learning may be easily adapted to address biological questions in an intuitive way. In particular, using this method we have identified a novel pluripotent state, which appears to be an intermediate between the well-known naïve and primed states (see Fig. 6) and shares many of the putative properties of a recently proposed ‘formative’ state (Smith, 2017). Cells in this state are characterized by partial dissolution of the core transcriptional regulatory circuit and distinct changes in cell-cell and cell-matrix interactions. It is unlikely that these cells correspond to the primed pluripotent state, since the culture conditions (low serum and LIF) in which ‘formative’ cells are observed in large numbers do not support FGF/Activin-dependent self-renewal of primed pluripotent EpiSCs (Brons et al., 2007; Tesar et al., 2007). Furthermore, these cells only appear at low frequency in 2i culture conditions and transiently during the early stages of cellular reprogramming of MEFs to pluripotency. Taken together these results suggest that this ‘formative’ state is a temporary intermediate in which the feedback mechanisms that stabilize the core pluripotency circuit become weakened and cells begin to become competent for lineage allocation. It remains to be seen how the population we have identified relates to recent observations of formative pluripotency characterized by loss of Rex1 expression and genome wide reorganization (Kalkan et al., 2017). We anticipate that the coming years will see greater advances in single cell profiling and analysis methods that will enable us to address this question, and identify with greater precision the regulatory networks that control the maintenance and exit from pluripotency.

## Materials and methods

### Single-cell expression data

Expression data from Zunder *et al*. (2015) (Zunder et al., 2015) was retrieved from the Cytobank repository (accession no. 43324). In summary, these data contain measurements of 46 features taken at the single-cell level by mass cytometry, from two separate engineered mouse embryonic stem cell (mESC) lines NG (Nanog-GFP) and NN (Nanog-Neomycin). Each mESC line contains doxycycline (dox) inducible gene cassettes for *Oct4, Sox2, Klf4* and *c-Myc* used for secondary reprogramming to pluripotency from somatic mouse embryonic fibroblasts (MEFs). Data includes the expression profiles of mESCs in steady state pluripotent stem cell culture conditions containing either Serum/LIF (denoted 0i) or Serum/LIF supplemented with 3*μ*M GSK3 inhibitor CHIR-99021 and 1*μ*M MEK inhibitor PD-0325901 (denoted 2i). Furthermore, time-course data comprised of snapshots of MEFs undergoing 16 days of dox treatment in MEF medium (DMEM, 10% serum) followed by 14 days without dox (123 medium + LIF) (Zunder et al., 2015). De-barcoded raw data was processed in R version 3.3.2 using the flowCore (Ellis et al., 2017) package version 1.40.4. Relevant features were logicle-transformed with parameters *w* = 0.6, *t* = 10,000 and m = 4.5.

### Cell cycle analysis

Classification of cell cycle status was performed based on the expression levels of Ki67 (absence indicates G0), phosphorylation of Histone H3 (presence indicates M) as described in Figure 4c of Zunder et al. (2015). Classification of G1-, G2- and S-phase was not possible due to a lack of discernible modes for marker IdU.

### Ensemble regulatory network

An ensemble model of binary node interactions (valid for an abstract average cell) was derived from publicly available data. Transcription factor binding data was derived from ChIPBase 2.0 (Zhou et al., 2017), and information on other known interactions were sourced from KEGG (Ogata et al., 1999) and Reactome.org (see **Table S1**).

## Statistical analysis

### Principal components analysis

Principal components analysis of scaled and centered training data (expression from mouse ES cells cultured in 0i conditions, see above) was conducted in R using the *prcomp* function.

### Gaussian mixture model

Gaussian mixture models were constructed in R using the *Mclust* package version 5.2.2 (Fraley and Raftery, 2002). Fit quality was assessed using the Bayesian information criterion (BIC). Minimum BIC indicates the best model fit, however, models with a higher number of parameters often only provide marginally better fits and the overall quality approaches a natural limit. Optimal trade off between increased parameters and quality of fit was obtained by selecting the model corresponding to the ‘elbow’ in the plot of fit quality against number of components.

### Density estimation

Estimate of the probability density function corresponding to the GMM identified above was obtained using the *densityMclust* function in R. Probability density estimates were calculated using the *predict* method in R.

### Classification

The GMM identified above was used for classification of data into either of four categories based on the highest posterior probability in combination with a reject option to avoid misclassification of vastly dissimilar phenotypes. Thus, points outside the 90^th^ percentile for all individual multivariate Gaussian distributions were rejected as outliers.

### Software and computer code

Analyses were performed in R version 3.3.2. Computer code used in this study is available as a R-markdown file from https://github.com/passt/Eigen-Networks.

### Data availability

Data used in this study is available from Cytobank (accession 43324).

## Author Information

### Acknowledgments

This research was funded by the Biotechnology and Biological Sciences Research Council, United Kingdom, grant number BB/L000512/1 and by the Medical Research Council, United Kingdom, grant number MC_PC_15078.

### Author contributions

Conceptualization, P.S.S. and B.D.M; Methodology and Investigation, P.S.S. and B.D.M.; Writing - Original Draft, P.S.S, Writing - Review & Editing, P.S.S., B.D.M.; Supervision, B.D.M.

### Competing interests

The authors declare no competing interests.

### Corresponding Author

Correspondence to P.S. Stumpf.

## Supporting information

**Supplementary Figure 1:**
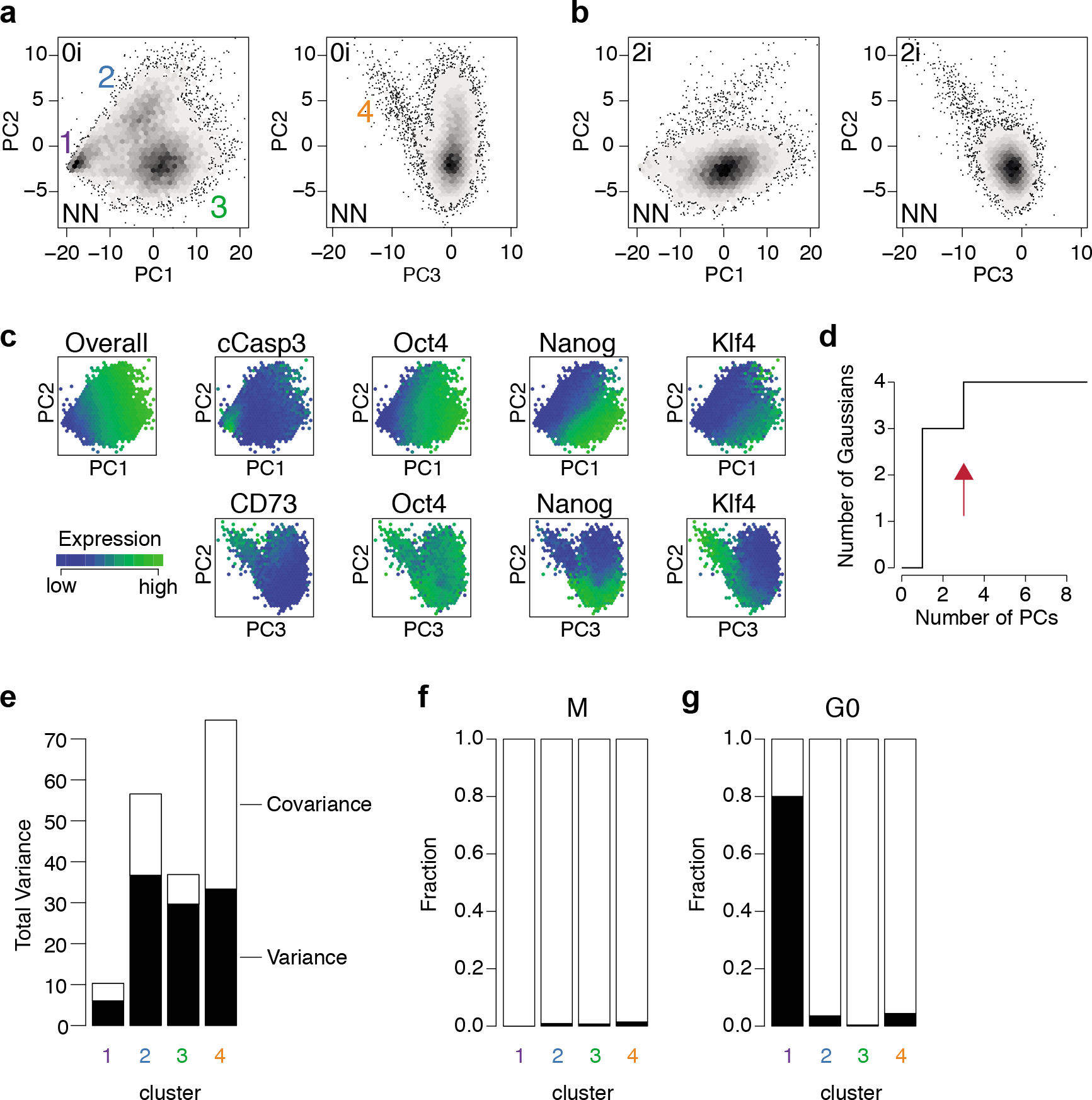
Sub-states of regulatory network activity. **(a-c)** Projection of Nanog-Neo (NN) mESC onto the same principal component space derived from Nanog-GFP (NG) mESCs (shown in Fig. 3). NN mESC display qualitatively the same population structure and corresponding node expression levels as NG mESCs. **(d)** Relationship between number of multivariate Gaussian distributions required to fully represent population structure, given the number of Principal Components used to represent network activity state. **(e)** Total variance/covariance within each sub-population (estimated from trace of the covariance matrix and the sum of the off diagonal elements of the covariance matrix for the respective fitted multivariate Gaussian models). **(f)** Fraction of cells of each cluster in M-phase of the cell cycle. **(g)** Fraction of cells of each cluster in G0-phase of the cell cycle.

